# The CRUNCH model does not account for load-dependent changes in visuospatial working memory in older adults: Evidence for the file-drawer problem in neuroimaging

**DOI:** 10.1101/451492

**Authors:** Sharna D Jamadar

**Affiliations:** Monash Institute for Cognitive and Clinical Neuroscience, Monash University. Wellington Rd, Melbourne, VIC 3800 AUSTRALIA; Monash Biomedical Imaging, Monash University. 770 Blackburn Rd, Melbourne, VIC 3800 AUSTRALIA; Australian Research Council Centre of Excellence for Integrative Brain Function. Melbourne, VIC 3800 AUSTRALIA

**Keywords:** compensation, cognitive ageing, CRUNCH model, selective reporting, reproducibility

## Abstract

Numerous neuroimaging studies have shown that older adults tend to activate the brain to a greater extent than younger adults during the performance of a task. This is typically interpreted as evidence for cognitive compensation. The Compensation-Related Utilisation of Neural Circuits Hypothesis (CRUNCH) model is a highly influential model of compensation, and states that increased functional magnetic resonance imaging (fMRI) activity in older adults compared to younger adults should reverse at higher levels of task difficulty. We tested the CRUNCH model using a visuospatial working memory paradigm, and found that fMRI activity in older vs. younger adults was in the opposite direction to that predicted by the model. Given that the CRUNCH model is the predominant model of compensation, this result was surprising. We followed up our results with a systematic review of the CRUNCH in healthy ageing literature using p-curve analysis. We find evidence for selective reporting, or the ‘file-drawer’ problem, in the cognitive compensation literature. Further experimental work is required to validate the CRUNCH model in cognitive ageing.

**Abbreviations:** CRUNCH: compensation-related utilisation of neural circuits hypothesis; fMRI: functional magnetic resonance imaging

**Highlights:**

- CRUNCH is the leading model of cognitive compensation in ageing
- We find fMRI activity in old vs. young adults in opposite direction predicted by CRUNCH
- We report quantitative evidence of selective reporting in CRUNCH literature

## 1. Introduction

A decline in cognitive function is considered an inevitable consequence of the ageing process. Even in healthy ageing adults, a small but definite decrease in most aspects of cognition is evident, particularly in processes that involve cognitive control or memory (Van Hooren et al., 2007). Despite well-documented decrements in cognitive function with age, and the potentially devastating effects of age-related decline on the individual’s quality of life, the mechanisms of age-related decline are not well understood.

There is substantial variation in the trajectory in which people age: some people show measurable decline early (in their 50s and 60s) whereas others continue on into their 70s, 80s and 90s showing little decline in their functional ability. One mechanism that may help individuals ameliorate age-related cognitive decline is the use of compensatory strategies. Biologically, the ageing process is accompanied by widespread changes in the functional and structural integrity of the brain (see Grady, 2012; Jamadar, 2018 for reviews), and it is commonly held that older adults can compensate for these changes by employing new strategies and/or additional brain regions to successfully perform cognitive tasks. One particularly robust effect found in many studies of cognitive ageing is that older adults often show greater activation in task-relevant networks, or activation of additional regions in comparison to young healthy adults. This additional brain activity is often associated with intact behavioural performance, leading many researchers to conclude that the additional activity is compensatory. Interestingly, similar effects have also been reported in psychiatric and neurodegenerative populations, including schizophrenia (Jamadar et al., 2010), Parkinson’s disease (van Nuenen et al., 2009), pre-manifest Huntington’s disease (Georgiou-Karistianis et al., 2013; Soloveva et al., 2018), and Alzheimer’s disease (Gould et al., 2006) among others; suggesting that compensation may be a prototypical mechanism for adapting to changes in the integrity of the brain.

For some time, the compensation hypothesis was an informal consensus that had arisen from numerous neuroimaging studies across a broad range of cognitive processes. It became axiomatic to link increased brain activity in an older (or psychiatric or neurodegenerative) population compared to young healthy adults to a compensatory process. This informal model of compensation was used to account for highly variable patterns of over-activity in experimental groups (Reuter-Lorenz and Cappell, 2008; Zarahn et al., 2007): while the phenomenon was initially used to account for increased brain activity that co-occurred with intact behavioural outcomes, many studies subsequently suggested that compensation may be at play even when performance was impaired in the experimental group. It was argued that even when over-activity is accompanied by performance decrements, it is likely that performance would be even worse without the additional activity – in other words, this pattern represented *failed* compensation. Thus, the informal model of compensation can account for *any* pattern of brain-behaviour changes, and so it is impossible to scientifically test or falsify the model. Furthermore, as it is not possible to falsify the informal model of compensation, it is also impossible to test it against competing models of brain-behaviour changes in ageing and pathology, such as dedifferentiation (Rajah and D’esposito, 2005).

The only formalised and falsifiable model of compensation proposed to date is the *compensation-related utilisation of neural circuits hypothesis* (CRUNCH; Reuter-Lorenz and Cappell, 2008). CRUNCH proposes that during task performance, as task difficulty (or load) increases, more cortical regions will be activated. Older adults reach their load capacity sooner than younger adults, so at easy and intermediate levels of task difficulty, they will recruit more neural resources than younger adults – the classic ‘compensation’ effect. At higher levels of load, the compensatory mechanism is no longer effective, leading to less activation and poorer performance in older vs. younger adults. Figure 1 shows the relationship between cognitive load and brain activity as proposed by CRUNCH. To test CRUNCH, it is necessary to use cognitive paradigms that parametrically manipulate four or more levels of load, because the predicted non-linear function requires at least four points to be described (Fabiani, 2012; Schneider-Garces et al., 2010). Evidence for CRUNCH has been obtained primarily in memory tasks. Using a verbal working memory task with 3 memory loads, Cappell et al. (2010) found a CRUNCH effect in right dorsolateral prefrontal cortex (BA46), with sub-threshold effects in BA9 of right dorsolateral prefrontal cortex, and right ventrolateral prefrontal cortex (BA45). Using a similar verbal working memory task, Schneider-Garces et al. (2010) found over-activity at low loads and under-activity at high loads in a fronto-parietal network in older compared to younger participants. Further support for CRUNCH was obtained in visuospatial working memory (Bauer et al., 2015; Toepper et al., 2014) and an n-back task (Mattay et al., 2006).

**Fig.1:**
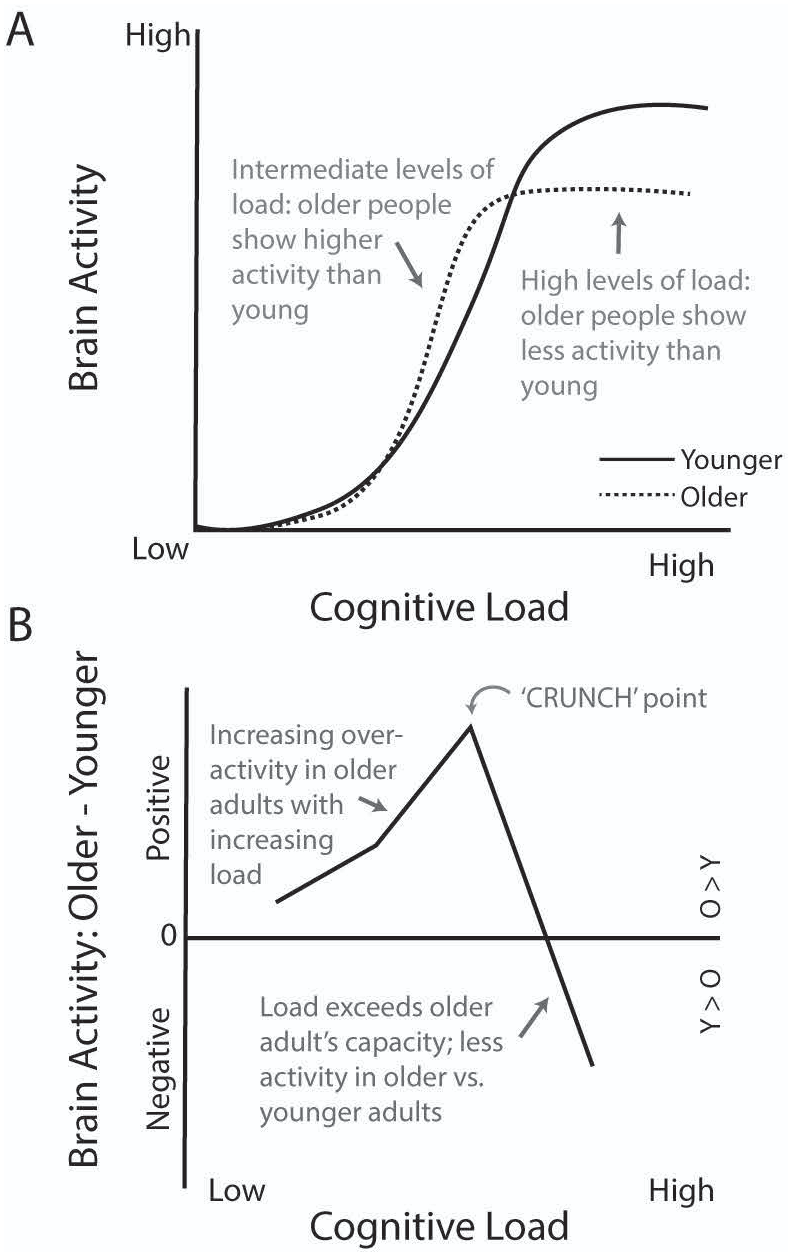
Hypothesised relationship between fMRI activity and cognitive load, as described in the CRUNCH model. (A) The non-linear relationship between fMRI activity and cognitive load requires at least 3 levels to be defined. At low or intermediate levels of load, older people show more activity than younger people. At high levels of load, older people exceed their processing capacity and show less fMRI activity than younger people. (B) The CRUNCH model is shown as the difference in fMRI activity between older and younger adults (older minus younger). The CRUNCH point is defined as the point where cognitive load exceeds older adults’ processing capacity, and older adults start to show less activity than younger adults. Note that according to the CRUNCH model, at low levels of load, older adults’ fMRI activity may match that of younger adults, or be slightly higher than younger adults.

In this study, we aimed to examine the links between cognitive compensation and quality of life outcomes. Age-related decrements in cognition have wide-ranging effects on the psychosocial outcomes of the individual, and their ability to carry out the day-to-day activities of life. While the CRUNCH model has established a quantitative metric of compensation using behavioural and fMRI data, little attention has been paid as to whether people who do compensate have better life outcomes. The current study was the first of a larger program of work that aimed to extend the CRUNCH model to cognitive domains beyond memory, and to examine links between the neural bases of compensation and psychosocial outcomes. Here, we aimed to confirm the CRUNCH effect in a working memory paradigm prior to extending it to a cognitive control (task-switching paradigm; results not reported here). We hypothesised that (a) behavioural results and fMRI activity in response to a visuospatial working memory task in older and younger adults would be consistent with the CRUNCH model; and (b) that the fMRI response to increasing task load would be positively associated with quality of life measures.

## 2. Study 1: Test of CRUNCH Predictions in Visuospatial Working Memory

### 2.1 Methods

#### 2.1.1 Participants

Forty-four individuals participated in this study. Four participants were excluded for not reaching a minimum performance threshold (see Procedure); two participants were excluded for suspected non-MR compatible implant; one participant was excluded for previously-unknown cerebral abnormality identified in the radiology report; and one person was excluded for suspected major depressive disorder. Data from 36 individuals were retained in the final sample, and demographics for the young and older groups are shown in Table 1.

**Table 1:**
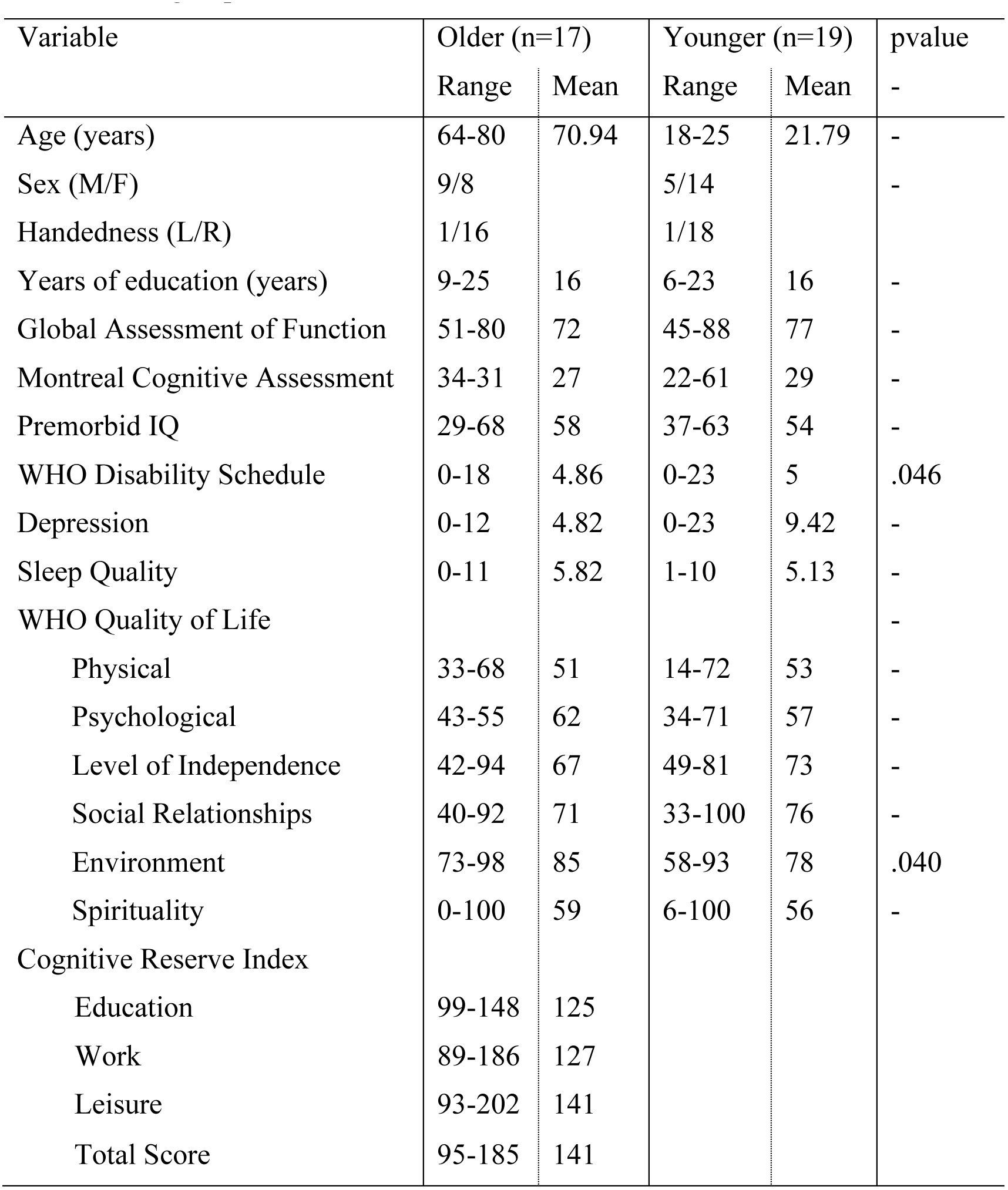
Demographic and psychosocial measures for each group. P-values indicate significant differences between each group.

Young and old groups did not differ in sex, handedness, years of education, global assessment of function (Hall, 1995), Montreal Cognitive Assessment (Nasreddine et al., 2005), Advanced Clinical Solutions Premorbid IQ (formerly Weschler Test of Adult Reading), Center for Epidemiological Studies Depression inventory-revised (CESD-R; Eaton et al., 2004), Pittsburgh Sleep Quality Index (PSQI; Buysse et al., 1989). The older group showed slightly reduced scores on the World Health Organisation Disability Schedule (WHODAS), which is an assessment of overall health and ability to carry out the activities of daily life. The younger group showed slightly lower ratings on the ‘environment’ subscale of the World Health Organisation Quality of Life-100 (WHOQOL) index, which reflects satisfaction with a person’s physical safety, home environment, financial resources, access to health care, social participation and transportation. In all other quality of life subscales, the groups did not differ.

Older adults also completed the Cognitive Reserve Index Questionnaire (CRIQ; Nucci et al., 2012). This assessment confirmed that our older group were well above the means calculated by Nucci et al.

Thus, overall the older and younger groups did not differ significantly in psychosocial function or general cognition.

#### 2.1.2 Stimuli and Tasks

The working memory task parametrically varied the working memory load across four conditions. Figure 2 shows an example trial sequence. The stimulus display was comprised of four boxes outlined in black, and labelled A, B, C, D. Two boxes were presented on the left (A, B), and two boxes on the right (C, D), with a fixation cross presented centrally. Participants responded with the index and middle fingers of their left and right hands. Each response mapped spatially with the stimulus display: A – left middle; B – left index; C – right index; D – right middle.

**Fig.2:**
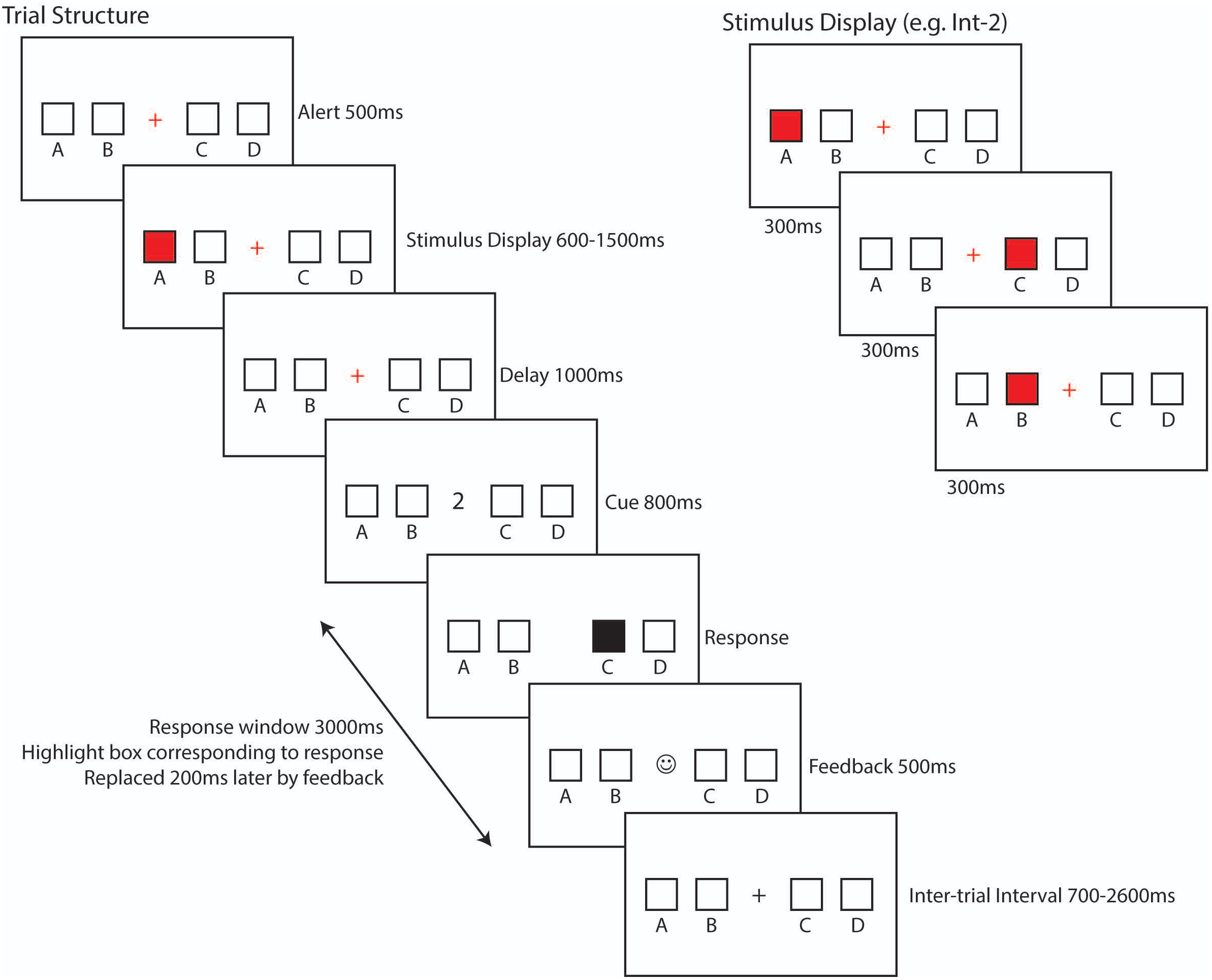
Visuospatial Working Memory paradigm. Participants viewed a screen with four boxes, which corresponded spatially with four responses (left middle, left index, right index, right middle). Following an alerting cue, the stimulus display was shown. The stimulus display showed either 2 consecutively highlighted boxes (low load), 3 (intermediate-1), 4 (intermediate-2) or 5 (high) highlighted boxes. Following a delay, participants were shown a cue (digit 1, 2, 3, 4, or 5) which indicated participants were required to respond with the box that was highlighted at that point in the sequence. In the example shown, the sequence was A, C, B, the cue was ‘2’, box C was highlighted second in the sequence, so the correct response was ‘C’. Feedback was shown for 500ms, then replaced with the fixation cue for the remainder of the trial.

At the start of each trial, the fixation cross was coloured red, to indicate that the next trial will start after a fixed duration of 500ms. The target display was then presented, where participants were presented with a sequence of red filled boxes, e.g., A, C, B. Each box in the sequence was highlighted for 300ms. After a 1000ms delay, the central fixation cross was replaced by a numerical cue; this indicated the position in the sequence that must be recalled on this trial. So, for a sequence of A, C, B, if the cue is ‘2’, then participants must indicate that ‘C’ was the second presented in the sequence, so the participant would press the button ‘C’ – right index finger. The participant’s response was indicated visually by a filled black box corresponding to their response, which was replaced 200ms later by feedback, where the central fixation cross was replaced by a tick or a cross for 500ms. The entire response window was 3000ms; for the remainder of the response window, the empty stimulus display with black fixation cross was shown until the beginning of the next trial.

Working memory load was varied by manipulating the number of highlighted boxes in each sequence. For Low load condition, two boxes were highlighted; for intermediate-1 load (int-1), 3 boxes were highlighted; intermediate-2 (int-2) 4 boxes, and high 5 boxes. The presentation of targets within each trial was pseudorandomised, so that each target position was equiprobable, there were no more than two instances of the same target location per trial, and no immediate repetitions of the same target location. The presentation of trials was also pseudorandomised, so that there were no more than four consecutive trials of the same load condition, no more than 3 consecutive cues indicating the same target position to be recalled, no more than 3 consecutive trials of the same response, and each response equiprobable. The total trial duration ranged from 5100ms (low) to 6000ms (high) depending on the current load.

#### 2.1.3 Procedure

All procedures were reviewed by the Monash University Human Research Ethics Committee, and complied with the Australian Code for the Responsible Conduct of Research (2007) and the Australian National Statement on Ethical Conduct in Human Research (2007).

Participants completed two testing sessions scheduled no more than two weeks apart. Approximately 1 week prior to their first session, psychosocial measures (WHOQOL, WHODAS, CESD-R, PSQI, IPAQ, Edinburgh Handedness) were posted to participants, to complete within their own time. Any uncompleted forms were completed in the time between testing sessions.

Testing session 1 comprised of cognitive testing and training on the MRI tasks. Participants completed the Cognitive Reserve Index Questionnaire (Nucci et al., 2012), Montreal Cognitive Assessment (Nasreddine et al., 2005), and Advanced Clinical Solutions Premorbid IQ with the researcher. Participants then completed a short training on two MRI tasks, the working memory task and a task-switching paradigm (not reported here). Participants were required to reach a performance threshold above chance to participate; three older and one younger participant were unable to reach the performance threshold for the task-switching paradigm, and so were not included in the final sample.

Training for the working memory task consisted of three blocks. In the first block, 10 trials of the low condition only were presented to familiarise participants with the task. In the second trial, 10 trials of the int-1 condition were presented. In the third block, 10 trials of int-2 condition, followed by 10 trials of the high load condition were presented. Thus, training session one included 40 trials overall, with 10 trials of each condition presented in a blocked sequence.

Testing session 2 involved a short training session to re-familiarise participants with the tasks, and MRI scanning. The brief training consisted of three blocks: block 1 comprised 5 trials of the low load condition. Block 2 comprised 5 trials of the int-1 condition, and block 3 comprised 5 trials of int-2 followed by 5 trials of high load condition. Thus, training session two included 20 trials overall, with 5 of each trial presented in a blocked sequence.

During the MRI scan, participants laid supine in the scanner bore, holding a response device in each hand. Visual stimuli were back-projected onto a screen positioned approximately 1m from the back of the bore. Participants viewed stimuli through a mirror mounted on the head coil. Participants completed two tasks within the MRI session, the working memory and task-switching tasks; the results of the task-switching paradigm are not reported here. Order of presentation of each task was counterbalanced between participants to control for fatigue. In the working memory task, participants completed three blocks of 43 trials, with proportionate numbers of each condition within each block. Overall, participants completed 31 trials each of low, int-1, int-2 conditions and 37 trials of high condition.

#### 2.1.4 Data Acquisition and Analysis

##### 2.1.4.1 Behavioural Data

Reaction time and accuracy (percent correct) data were analysed with separate 2 group (young, old) x 4 difficulty (low, int-1, int-2, high) repeated measures ANOVAs.

##### 2.1.4.2 MRI Data

Magnetic resonance images were acquired on a Siemens Skyra 3T scanner using a 32 channel head coil. Functional MRI was acquired using a T2*-weighted GRAPPA echo-planar imaging (EPI) sequence (ascending axial acquisition, TR = 2.5s, TE = 30s, FOV = 192mm, acquisition matrix = 64 x 64, 44 slices, 3 x 3 x 3mm voxels, 150 volumes per block). Structural MRI was acquired using a T1-weighted 3D MPRAGE sequence (TR=1900ms, TE = 2.43ms, flip angle = 9°, FOV = 192 x 192mm, voxel size = 0.6mm, 256 slices).

MRI data were analysed with SPM12 (Wellcome Department of Cognitive Neurology, London). For functional runs, the first five images were discarded to account for T1 saturation effects. EPIs were realigned to the first image in each run and co-registered to each individual’s T1 structural scan. T1 scans were then segmented using the unified segmentation algorithm in SPM12 to derive parameters to normalise from individual subject space to MNI space. Functional and structural scans were then normalised to the MNI template using these parameters and spatially smoothed using a 6 x 6 x 6mm FWHM Gaussian kernel. Quality of registration was checked and realignment parameters screened for motion greater than 1mm.

Events for each participant were categorised as correct or incorrect, for each condition separately. On average, participants generated 29 correct low trials, 28 correct int-1 trials, 24 correct int-2 trials and 25 correct high load trials. First-level analyses consisted of a model with the eight experimental conditions (correct low, correct int-1, correct int-2, correct high; incorrect low, incorrect int-1, incorrect int-2, incorrect high), and six realignment parameters and convolved with a canonical haemodynamic response function with temporal derivative. Due to the small trial numbers for incorrect trials, these were not further analysed.

To determine if the groups differed in grey matter volumes, voxel based morphometry analysis (Ashburner and Friston, 2000), implemented in cat12 toolbox (Gaser and Dahnke, 2016), was performed. The independent samples t-test controlling for total intracranial volume confirmed that the older group showed smaller grey matter volumes across the brain (maximum t(33)= 9.09, p<.001). Thus, total grey matter volume was included as a covariate at the second-level.

At the second-level, a 2 group x 4 difficulty full-factorial ANOVA controlling for grey matter volume was conducted and corrected for multiple comparisons at a whole-brain level at p<.001, FWE corrected at the cluster level. Region-of-interest (ROI) analyses were conducted to determine the direction of any significant effects; details of the ROI analyses are given in the Results section.

### 2.3 Results

#### 2.3.1 Behavioural Results

RT and percent error are shown in Figure 3. Overall, older adults were slower than younger adults (F(1,34)=14.63, p=.001) and reaction time increased linearly with difficulty (F(3,102)=90.47, p<.001; linear effect F(1,34)=164.49, p<.001). The effect of difficulty did not differ between the groups (p=.649).

**Fig.3:**
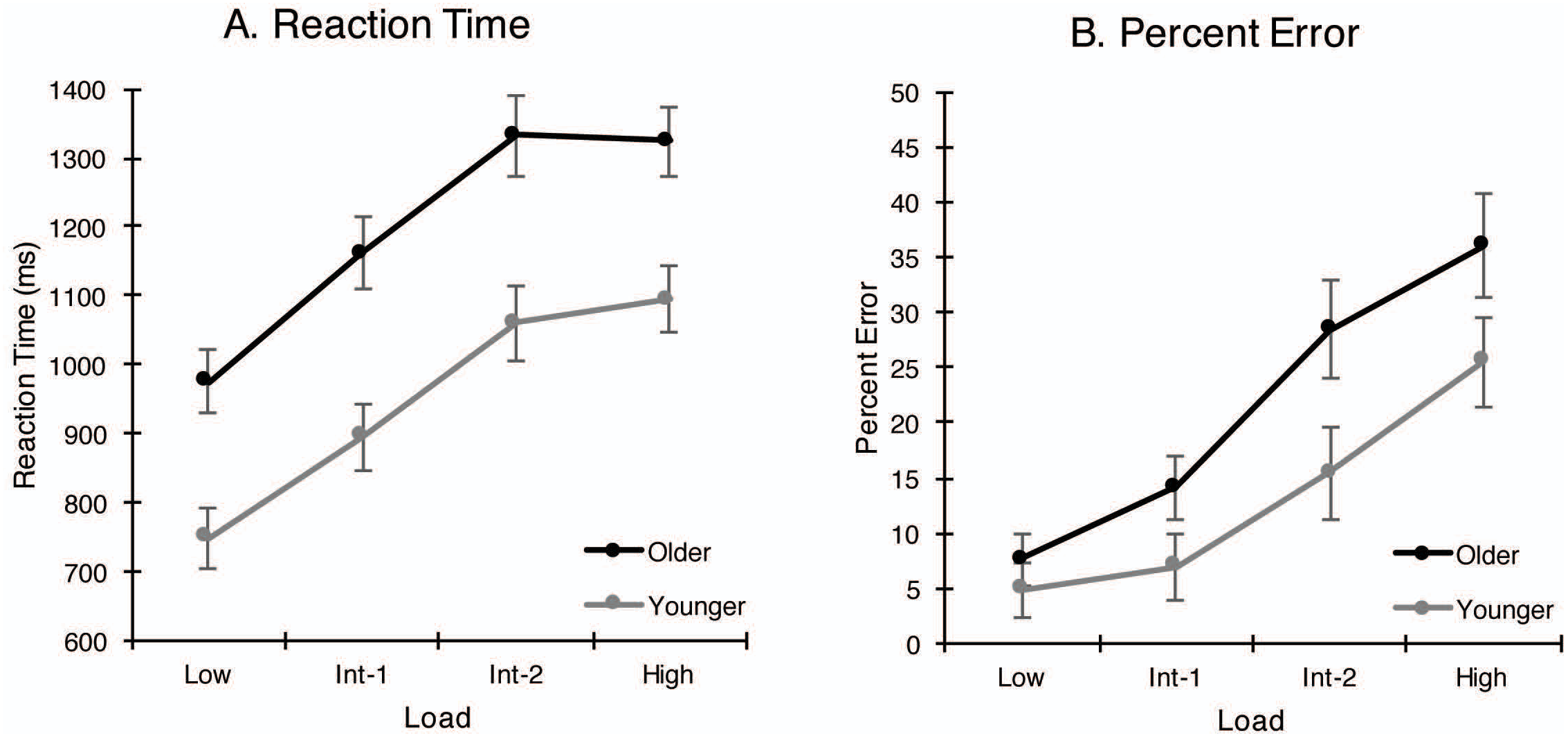
Behavioural results. (A) Reaction time (ms); (B) Percent error. Error bars show standard error. Abbreviations: Int-1, intermediate-1; Int-2, intermediate-2.

Older adults were less accurate than younger adults (F(1,34)=7.514, p=.010), and accuracy decreased linearly with increased difficulty (F(3,102)=24.30, p<.001). The group x difficulty interaction (F(3,102)=8.39, p=.001) confirmed that the reduction in accuracy with increased difficulty was larger for older adults than younger adults.

#### 2.3.2 MRI Results

The main effect of group must be interpreted with caution, given that group differences in haemodynamic response function were not controlled for. However, for completeness, we report this effect here. After controlling for grey matter volume, the main effect of group revealed a single cluster within the right caudate (F=26.21, k=80 voxels, MNI: 18, −2, 22). Parameter estimates were examined to determine the direction of the effect. A sphere of diameter 10mm was created around the voxel of maximum intensity within the cluster, and parameter estimates (beta values) were extracted from the ROI using MarsBar (v0.44, Brett et al., 2002). The ROI analysis confirmed that older adults (M=1.02, SE=0.32) showed larger activity in the right caudate than younger adults (M=−0.76, SE=0.43).

The main effect of difficulty revealed widespread effects throughout the brain at this statistical threshold (Table 2; Figure 4). Parameter estimates were examined to determine the direction of the effect. To constrain the ROI analysis, anatomical masks of major prefrontal, subcortical and parietal subdivisions were created using the aal definitions (Tzourio-Mazoyer et al., 2002) implemented in wfu_PickAtlas tool (v2.4, Maldjian et al., 2003). These masks were then applied to the F-map for the main effect of difficulty; within each region, a sphere (diameter 10mm) was created around the voxel of peak intensity, and parameter estimates were extracted from the region using MarsBar (Brett et al., 2002). Most (15/22) regions showed a roughly positive linear relationship between BOLD activity and task difficulty (Table 2), consistent with the design of the paradigm. Five additional regions showed a roughly inverted-U shaped effect of difficulty, with trend of increasing activity for Low-Int1-Int2 loads, and a decrease in activity between Int2-High loads. The remaining 2 regions (left anterior cingulate, right putamen) showed effects of difficulty that were not consistent with the design.

**Fig.4:**
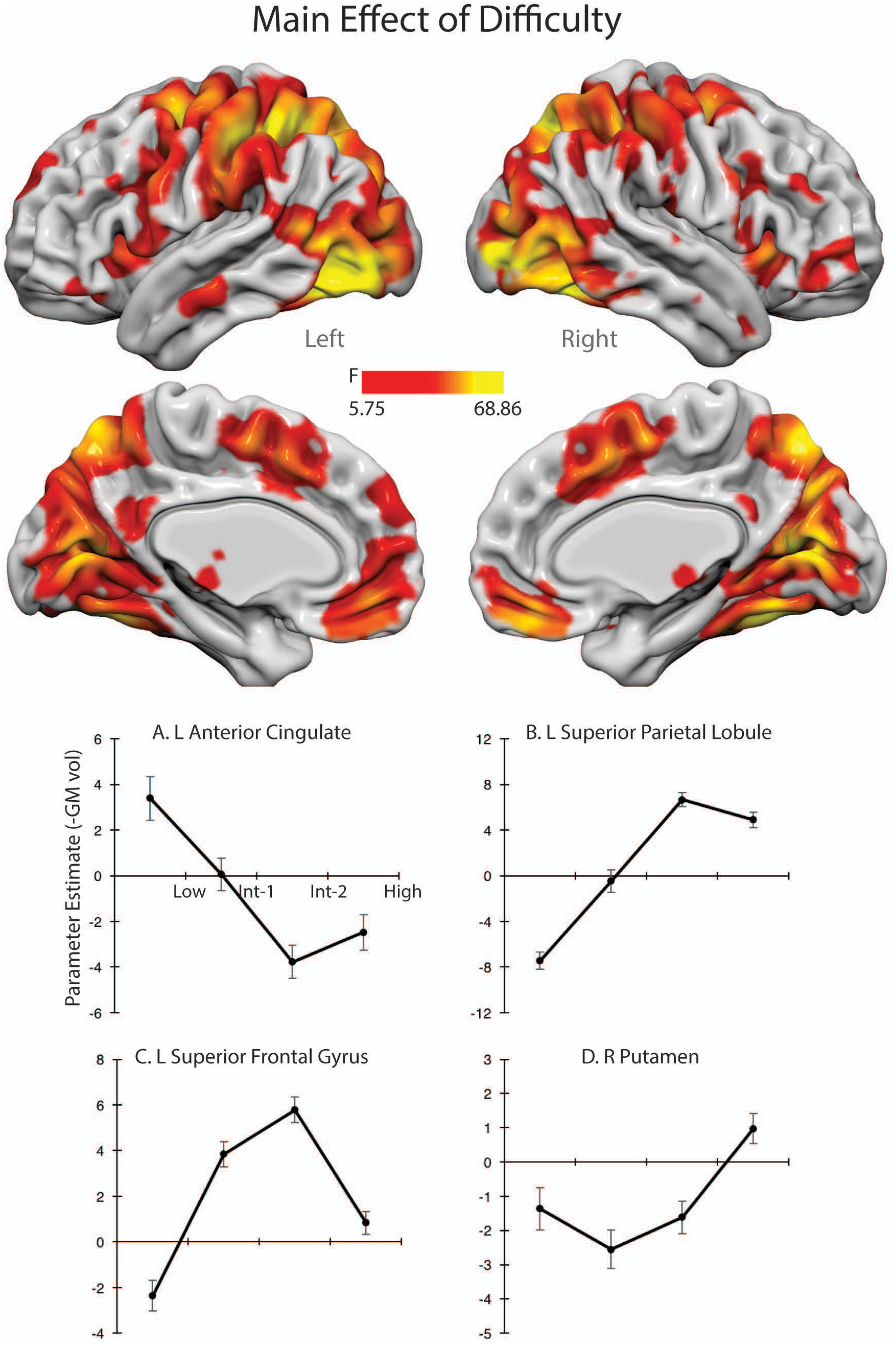
fMRI results for the main effect of difficulty. Plots show four representative effects of load (Values presented in Table 1). (A) Left anterior cingulate showed a negative linear effect of load. Note that no other region showed a negative linear effect of load. (B) Left superior parietal lobule showed a positive linear effect of load; 14 (out of 22) other regions showed this pattern of results. (C) Left superior frontal gyrus showed a negative quadratic effect of load; 5 (out of 22) other regions showed this pattern of results. (D) Right putamen showed a positive quadratic effect of load; 1 (out of 22) other region showed this pattern of results. Error bars show standard error. Abbreviations: L, left; R, right; Int-1, intermediate-1; Int-2, intermediate-2.

**Table 2:**
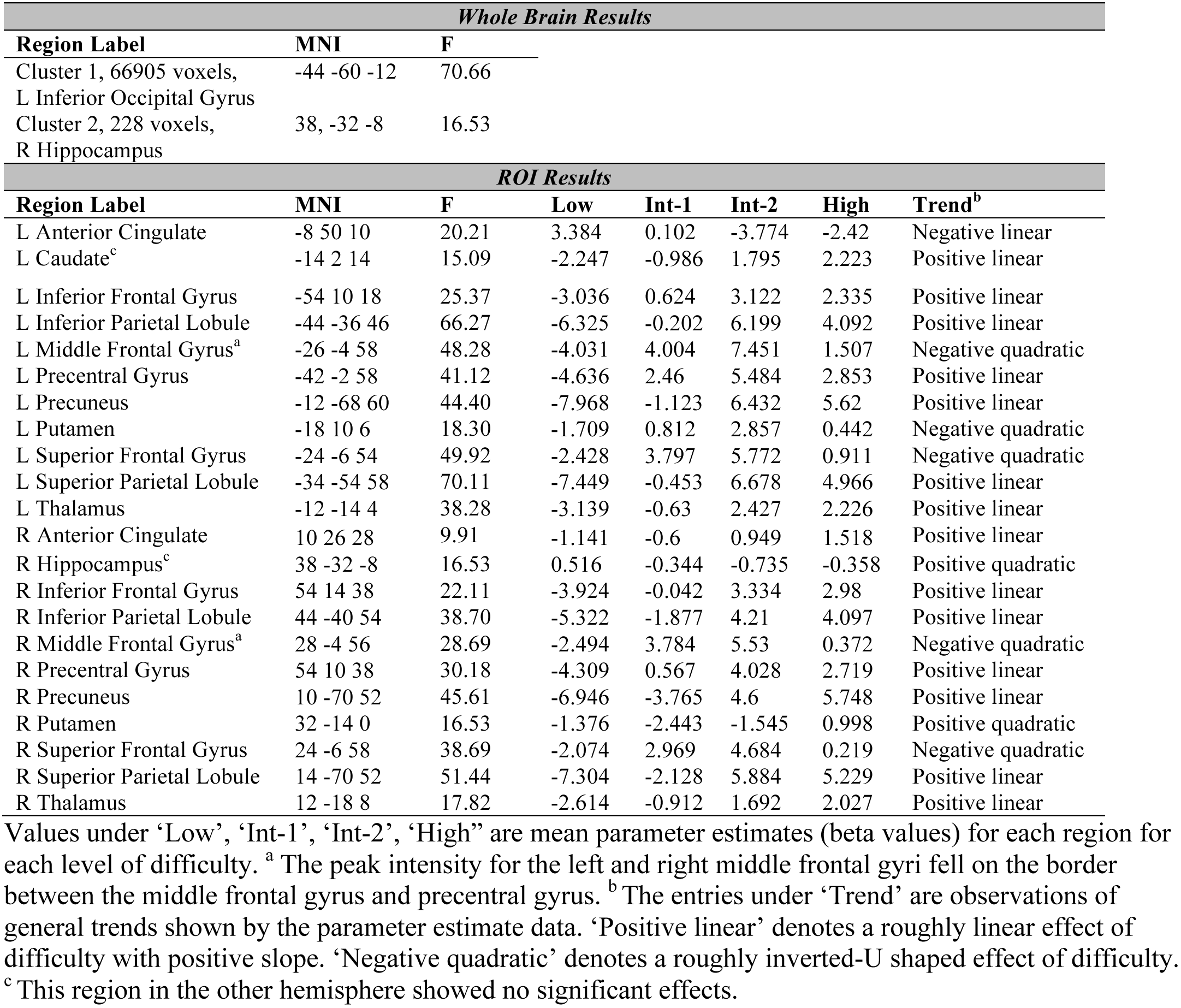
MNI coordinates and F values for regions showing significant effects of difficulty.

The group x difficulty interaction, controlling for grey matter volume, revealed significant effects in ventrolateral prefrontal cortex, premotor cortices and subcortical regions (Table 3; Figure 5). To determine the direction of the effect, an ROI analysis was conducted by creating a sphere (diameter 10mm) around the peak intensity voxel within each cluster; parameter estimates were extracted using MarsBar. Figure 5 shows the parameter estimates for the group x difficulty interaction. As a general trend, across regions fMRI activity increased with task difficulty, although this trend did not hold for all regions, and was not linear. At low levels of difficulty, there was a trend towards larger fMRI activity for younger vs. older adults, particularly in the left and right inferior frontal gyrus, left middle temporal gyrus, and left putamen. In the left precentral gyrus, older adults showed larger activity than younger adults at low levels of difficulty. At intermediate levels of difficulty, most regions showed no or small differences between the groups, the exception was the left precentral gyrus and left supplementary motor area, where younger adults showed larger activity compared to older adults. At the highest difficulty level, older adults showed greater activity than younger adults across regions.

**Fig.5:**
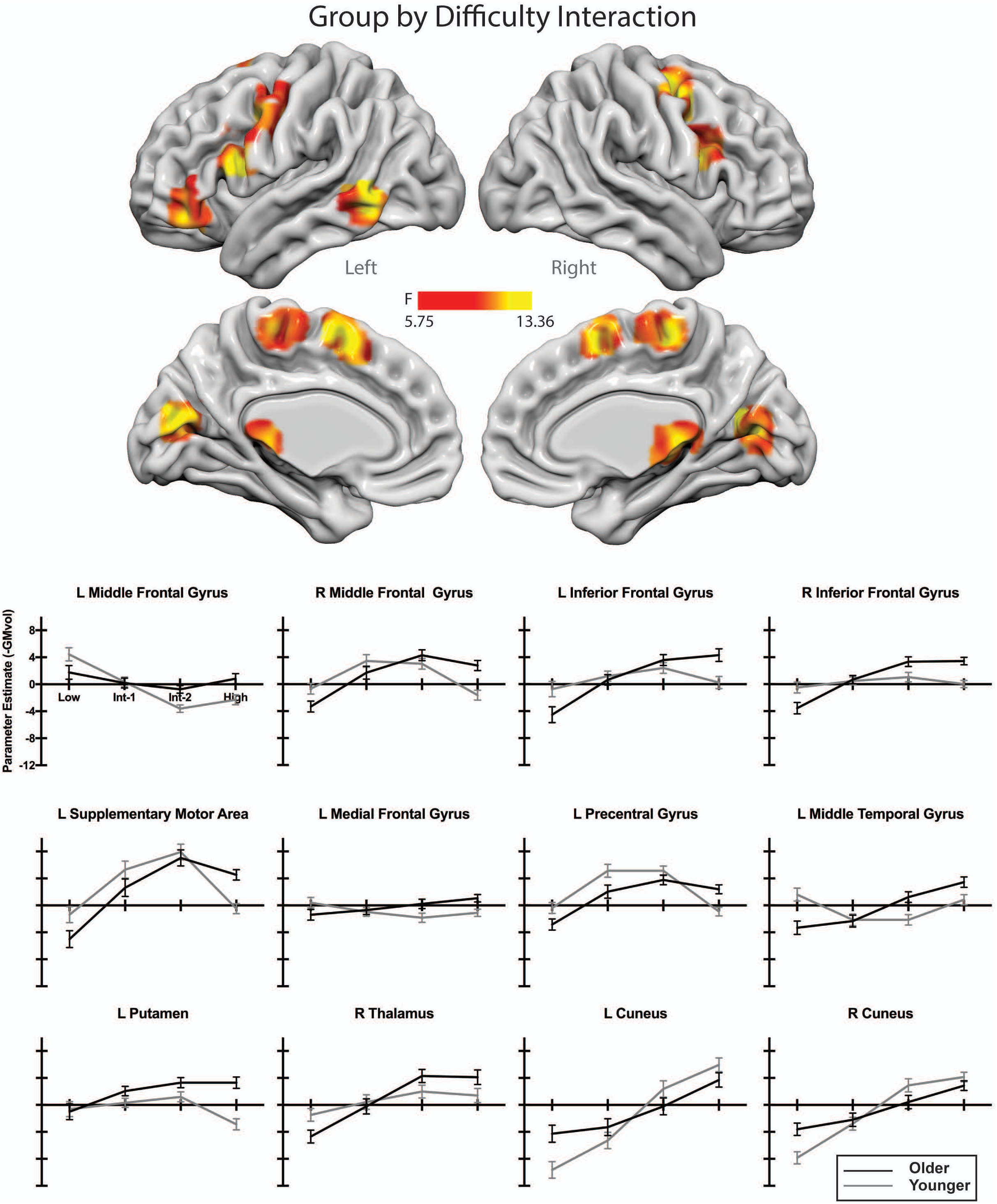
fMRI results for the group by difficulty interaction. Plots show the interaction for each region of interest. Error bars show standard error. Abbreviations: L, left; R, right; Int-1, intermediate-1; Int-2, intermediate-2.

**Table 3:** MNI coordinates and F values for regions showing a significant Group by Difficulty Interaction

| *Whole Brain Results* |  |  | *Simple Effects* |  |
| --- | --- | --- | --- | --- |
| **Region Label** | **MNI** | **F** | **Low Load** | **High Load** |
| Cluster 1, 84 voxels<br>L Middle Frontal Gyrus<br>L Inferior Frontal<br>Gyrus | -40 40 -10 | 8.532 | t(20)=-1.943 | t(31)=2.983 |
| Cluster 2, 105 voxels<br>L Middle Temporal<br>Gyrus<br>L Inferior Temporal<br>Gyrus | -60 -54 -6 | 9.410 | t(28)=-3.583 | t(23)=2.45 |
| Cluster 3, 103 voxels<br>R Thalamus | 8 -24 -2 | 9.333 | t(28)=-2.441 | t(27)=1.781 |
| Cluster 4, 183 voxels<br>L Putamen<br>L Caudate | -18 14 6 | 13.362 | t(23)=-.246 | t(32)=5.213 |
| Cluster 5, 86 voxels<br>R Cuneus<br>R Posterior Cingulate | 16 -70 14 | 8.848 | t(32)=3.259 | t(29)=-1.305 |
| Cluster 6, 82 voxels<br>L Cuneus | -8 -78 16 | 10.077 | t(32)=3.027 | t(22)=-1.53 |
| Cluster 7, 111 voxels<br>L Inferior Frontal<br>Gyrus | -52 12 18 | 11.361 | t(28)=-2.33 | t(32)=3.119 |
| Cluster 8, 142 voxels<br>R Inferior Frontal<br>Gyrus | 46 10 22 | 8.997 | t(23)=-2.655 | t(21)=4.342 |
| Cluster 9, 137 voxels<br>L Precentral Gyrus | -42 -4 44 | 11.660 | t(25)=-2.154 | t(32)=3.543 |
| Cluster 10, 212 voxels<br>R Middle Frontal Gyrus<br>R Precentral Gyrus<br>R Superior Frontal<br>Gyrus | 36 -4 52 | 12.563 | t(23)=-2.377 | t(32)=4.179 |
| Cluster 11, 128 voxels<br>L Supplementary<br>Motor Area<br>R Supplementary<br>Motor Area | -2 10 56 | 10.174 | t(28)=-2.122 | t(32)=4.73 |
| Cluster 12, 79 voxels<br>L Medial Frontal Gyrus | 6 -30 64 | 9.002 | t(24)=-1.649 | t(27)=2.833 |

Simple effect of group at low and high load levels tested using ROI data, independent samples t-test, equal variances not assumed. Abbreviations: L, left; R, right.

To visualise the group x difficulty interaction, Figure 6 shows older-younger difference scores for each level of difficulty; the top left panel (Figure 6A, cross marks) shows an example of predicted group differences at the critical levels of task difficulty (low and high loads) as hypothesised by CRUNCH. As can be seen from Figure 6, all regions except the cuneus showed the opposite relationship between group and difficulty level as predicted by CRUNCH. The bilateral cuneus showed greater activity in older adults than younger adults at low loads, and less activity in older than younger adults at high loads; consistent with the predictions of the CRUNCH model. To test if these effects were significant, we ran *post hoc* independent samples t-tests at low and high levels on parameter estimates calculated from the group x difficulty for the cuneus. These *post hoc* tests were controlled for multiple comparisons (4 tests: left & right cuneus, low and high loads; α =0.05/4; p=0.0125) confirmed that the group difference was significant at low loads (left cuneus: t(33)=3.027, p=.005; right cuneus, t(33)=3.259, p=.003) but not at high loads (both p>.138).

**Fig.6:**
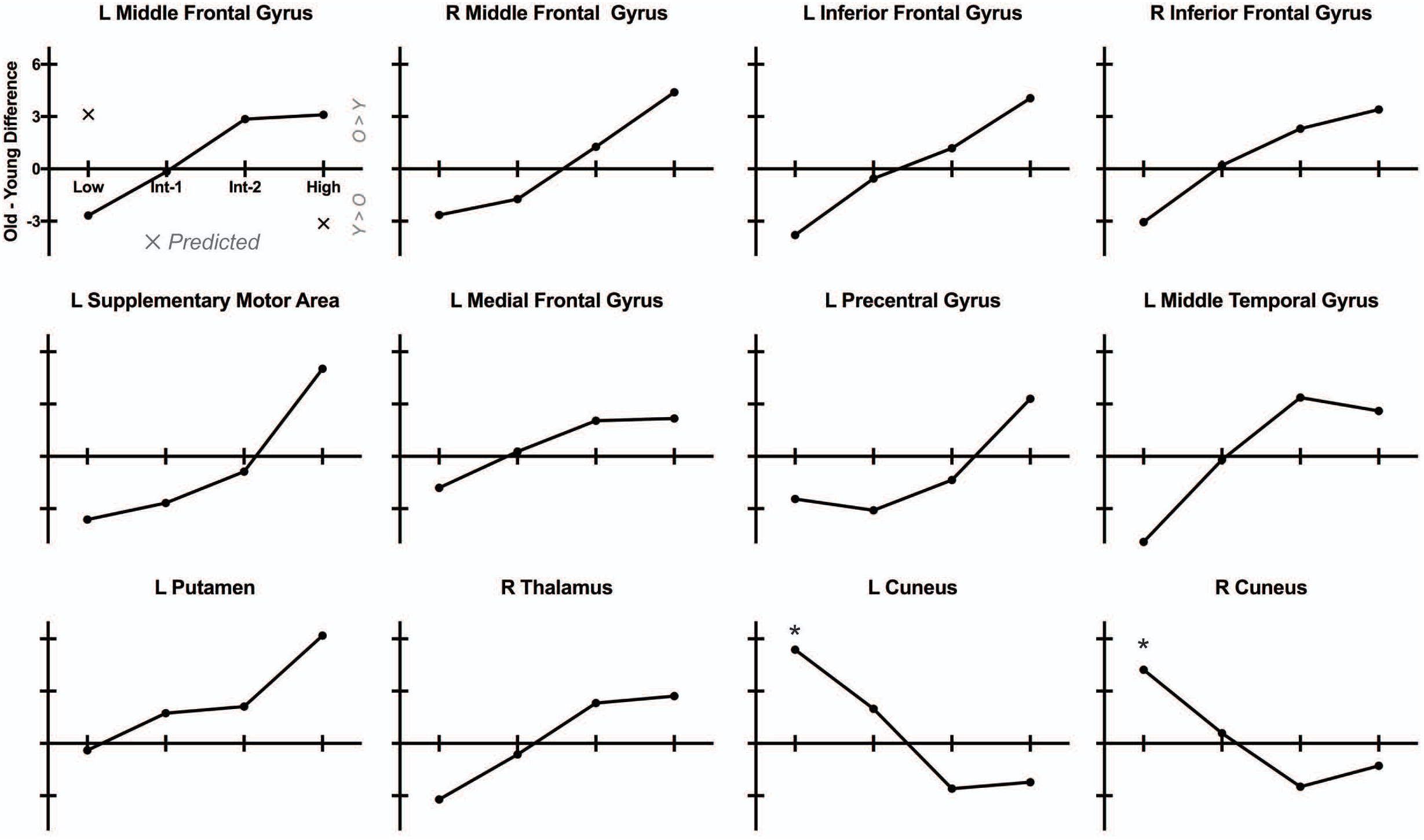
Older minus younger parameter estimate differences for each region showing a significant group by difficulty interaction (see Fig.5). Positive values indicate that older adults showed greater activity in this region compared to younger adults. Negative values indicate that younger adults showed greater activity than older adults. Panel A shows predicted difference values (indicated by cross markers) at the critical load levels (low and high) according to CRUNCH. According to the model, older adults should show greater activity than younger adults at low loads, and the opposite effect at high loads. Abbreviations: L, left; R, right; Int-1, intermediate-1; Int-2, intermediate-2.

#### 2.3.3 Relationship between brain activity and psychosocial function

A primary goal of this research was to examine the relationship between the neural CRUNCH effect and psychosocial outcomes. To this end, for each ROI showing a rough CRUNCH effect (left and right cuneus), we estimated beta scores for the linear regression between difficulty (independent) and ROI BOLD activity (dependent) for each individual. Betas for each individual was submitted to Pearson bivariate correlations with WHODAS and WHOQOL overall scores. Correlations were corrected for multiple comparisons for each ROI (α=0.05/2, p=0.025). In the older group, the CRUNCH effect was not associated with WHODAS or WHOQOL (all p >.202). In the younger group, the CRUNCH effect was positively associated with WHODAS (both r=.488, p=0.04), however this did not survive correction for multiple comparisons.

### 2.4 Discussion

In this study, we aimed to examine the links between the CRUNCH effect during working memory and psychosocial outcomes. We used a novel task that varied visuospatial working memory load from 2 items (low) to 5 items (high load). The behavioural and fMRI results confirmed that the task was successful in manipulating task difficulty. RT and error rate increased linearly with increasing task load. Despite overall elevated RTs and error rates, older adults showed an effect of load that was not significantly different from younger adults, consistent with the conclusion that older adults showed intact task performance. At the neural level, most but not all regions (15/22) showing a main effect of task difficulty showed a positive linear effect, with fMRI activity increasing with increasing task difficulty. So, at the behavioural and neural level, the novel visuospatial working memory task was successful in (a) manipulating task difficulty across four levels of load, and (b) showing intact behavioural performance for the older group; as required to test the CRUNCH model (Fabiani, 2012; Schneider-Garces et al., 2010).

The crucial test of the CRUNCH model, the fMRI group by difficulty interaction, showed little consistency with the predictions of the model. We obtained a significant group by difficulty interaction in a distributed network encompassing prefrontal, premotor, subcortical and visual regions. The group by difficulty interaction in all regions except the bilateral cuneus showed an opposite effect to that predicted by the model (Figure 6); with prefrontal, premotor and subcortical regions showing an increase in fMRI activity difference between the group with task difficulty. By contrast, the CRUNCH model predicts that at the group difference should increase until the highest loads, until the older adults reach ‘CRUNCH’ point, after which the group difference reverses. As such, our results in general did not support the model. The one exception to this pattern was the cuneus, which showed the expected group difference at low loads (old > young); and an effect of difficulty that was compatible with the CRUNCH predictions. The group difference at high loads (young > old) was not significant.

These results appear to be the first that do not support the CRUNCH model. To our knowledge, this is the first published paper reporting results inconsistent with the CRUNCH model. It could be argued that our highest level of difficulty was not sufficiently difficult to induce a CRUNCH effect. However, it is notable that our percent error for the high load condition in the older group (mean ∼35 %) was consistent with, or higher than error rates in previous tests of CRUNCH (e.g., Cappell et al., 2010; older percent error at high load ∼25%; Mattay et al., 2006, ∼20%; see Supplement, Study Details Table). Thus, we argue that the task was difficult enough to push older adults past their ‘CRUNCH’ point. It could also be argued that our sample size was insufficient to detect this effect. However, it is notable that while our sample size was modest, it is consistent with that of previous CRUNCH studies (e.g., Cappell et al., n=21 young, n=23 older; Mattay et al., n=10 younger, n=12 older; Bauer et al., 2015, n=19 younger, n=21 older; also see Supplement; Study Details Table). Furthermore, the results in all ROIs except the visual cortex showed effects in the opposite direction to that predicted by CRUNCH, suggesting that our modest sample size does not explain our failure to replicate the CRUNCH effect.

It is possible that our sample was not sufficiently impaired to require compensation. In favour of this argument are the results showing that the older group showed a high level of cognitive reserve as tested using the Cognitive Reserve Index Questionnaire (Table 1) and the groups did not significantly differ on Global Assessment of Function, Montreal Cognitive Assessment, or Quality of Life. However, in contrast to the argument that our older group was not sufficiently impaired to detect compensation, are the results showing that (a) the older group showed reduced grey matter volumes compared to younger adults; (b) the older group showed overall psychomotor slowing and elevated error rate compared to younger adults, consistent with cognitive ageing; (c) groups showed a marginally reduced ability to carry out day-to-day activities, as assessed using the WHO Disability Schedule; and (d) although direct comparison with previous studies are difficult given different methods, our older group appears to be similar to previous samples in terms of cognitive reserve (years of education) and current cognition (see Supplement; Study Details Table). Thus, although it remains possible that our sample was not sufficiently impaired to detect compensatory activity, our sample is comparable with previous samples that have detected compensation using CRUNCH.

It is notable that although the CRUNCH model remains highly influential in the literature, only a modest number of studies have directly tested its hypotheses, and the existing studies are restricted only to a single cognitive domain – memory. Other studies that have reported results consistent with the CRUNCH model have used methods that are difficult to compare to predictions based on fMRI activity (e.g., MEG, Proskovec et al., 2016; behavour only, Fu et al., 2017). This raises the question of whether publication bias is a factor in this literature, where only a small number of studies exist that explicitly test the model predictions. In order to examine if this is the case, we conducted a meta-analysis using p-curves (Simonsohn et al., 2014) of the CRUNCH literature.

## 3. Study 2: Systematic Review of CRUNCH Literature

### 3.1 Methods

The search for studies that test CRUNCH predictions was conducted systematically, in accordance with the protocol presented by Simonsohn et al. (2014) and the procedures given on their website (p-curve.com; v.4.06).

#### 3.1.1 Study Selection Rule

The study selection rule was all fMRI or PET studies that tested the CRUNCH model with at least 3 levels of task difficulty, and reporting a group x difficulty interaction.

Definition of CRUNCH model predictions were based on the report of Reuter-Lorenz & Cappell (2008). Specifically, behavioural predictions are based on accuracy: “According to CRUNCH, compensatory recruitment *at low demand maintains seniors’ performance at levels that are equivalent to or minimally different from younger adult levels*. [Reference to Figure not presented here] …*as task demands increase, older adults reach a resource ceiling, and performance levels drop, especially in comparison to those of younger groups*. At peak levels of demand, errors may be sufficiently frequent that the task is met with frustration or approached with ineffective strategies, or other factors may prevail that lead to underactivation of this region compared to younger groups” (page 180). With regards to brain activity: “Increased recruitment in response to increasing task demand is a ‘‘normal’’ neural response, evident in younger adults; what varies with age, according to CRUNCH, is the slope of the function relating activation to demand and the level at which activation asymptotes. [Reference to Figure not presented here] … relative to younger adults, *older adults progress from over-activation at lower levels of task demand to under-activation at higher levels of task demand within the same region of interest*.” (page 180).

The p-curve test is critically dependent on the accurate selection of the result of interest, and Simonsohn et al. (2014) provide guidelines for how to select p values for a range of experimental designs. On the basis of the CRUNCH predictions, we set the p-curve test of interest to a ‘3×2 design attenuated trend’ for accuracy data and ‘3×2 design reversing trends’ for fMRI ROI data.

#### 3.1.2 Manuscript Identification

Manuscripts were reviewed using a systematic approach consistent with the p-curve guidelines (Simonsohn et al., 2014; Simonsohn et al., 2015). Full report of the identified manuscripts and criteria for exclusion is given in the Supplement (Exclusion Table).

(1) We conducted a search on Google Scholar (July 2018) using the terms [(“compensation related utilization of neural circuits”) AND (“fMRI”) OR (“PET”)] and restricted the results to those published since 2008 (when the first CRUNCH paper was published). The search was also conducted on PubMed and Web of Science, however these searches only revealed 14 manuscripts each; the Google Scholar search was retained for the analysis as it was provided the highest hit rate. This resulted in 283 results.
(2) We then scanned the abstracts, and included only peer-reviewed experimental articles, excluding reviews, meta-analyses, conference abstracts, theses, papers that reported non-fMRI or PET experiments, and papers published in languages other than English. This yielded 52 results.
(3) Methods were reviewed to determine if cognitive load was varied across 3 or more levels, yielding 24 studies.
(4) Methods were reviewed to include only studies that compare young and old subjects, excluding ‘old-only’ or ‘middle-aged only’ samples; yielding 14 studies.
(5) Manuscripts were reviewed to exclude cases where the same participants contributed to more than one paper; two studies were excluded yielding 12 studies. Note that for some studies it was unclear if participants contributed data to more than one paper, and this is noted in the Supplement.

#### 3.1.3 P-Curve Disclosure Table

Central to the p-curve analysis is the comprehensive review of manuscripts and creation of the P-Curve Disclosure Table (see Supplement, P-Curve Disclosure Table). This Table summarises the stated aims and predictions, statistical tests and reported results. During the course of completing the Table, four studies were identified that did not report the critical group x load interaction to test the CRUNCH model, and these were removed from the analysis, leaving 8 studies that met criteria for inclusion in the p-curve analysis.

### 3.2 Results

The CRUNCH model predicts an interaction between age group and task difficulty, such that the difference between groups is small at low levels of difficulty, and large at high levels of difficulty. This effect is described in p-curve as an ‘attenuated’ interaction, and requires testing of the main interaction effect (Simonsohn et al., 2014). All 9 studies reported the interaction between age group and difficulty for accuracy data, and the pcurve is shown in Figure 7. The p-curve for accuracy data showed a strong right bias, suggesting that there is evidential value for the predicted accuracy age x difficulty interaction (full p-curve evidential value Z=-4.86, p<.0001; half-p-curve evidential value Z=-4.37, p<.0001).

**Fig.7:**
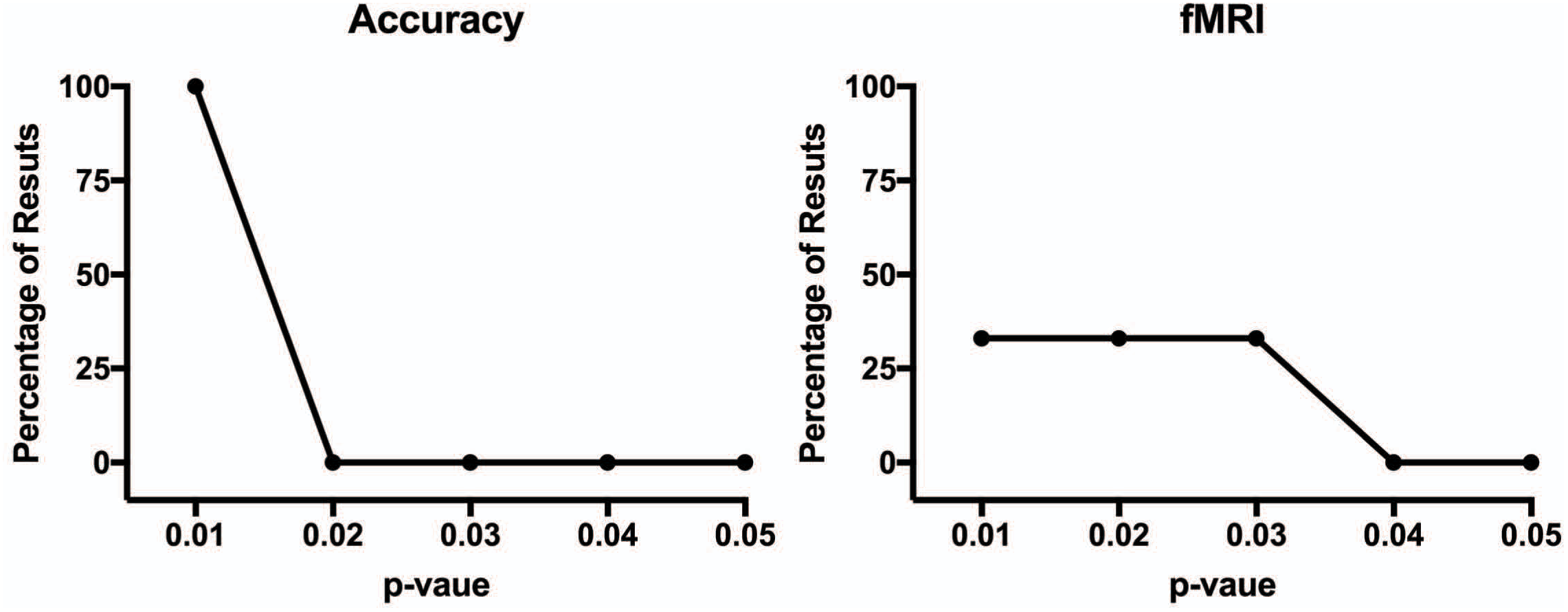
P-curve analysis for CRUNCH effects for (left) accuracy data and (right) fMRI data.

For fMRI, the CRUNCH model predicts a reversing trends interaction between age group and difficulty, such that the difference between young and old groups reverses at the extreme values of difficulty. This effect requires testing the age simple effect at the low and high levels of difficulty. Four of the 9 papers reported the required statistical tests. P-curve requires the values entered into the analysis to be independent, which is often not the case in fMRI studies, where results are repeated across the brain or multiple regions-of-interest. Therefore, to achieve the fairest and most unbiased estimate of the p-curve as possible, we firstly reviewed the magnitude of the age x difficulty interaction for each reported region. We identified the region that showed the strongest interaction effect, and then entered the simple effect results for that region into the p-curve analysis (see P-Curve Disclosure Table for values entered into the analysis for each paper). The p-curve for the strongest interaction effect for each manuscript is shown in Figure 7; in this case, the p-curve shows a flat distribution with slight right-ward bias. So, the p-curve suggests that there is not evidential value for the predicted fMRI age x difficulty interaction (full p-curve evidential value: Z=-1.71, p=.0437; half p-curve evidential value Z=-.029, p=.3877).

An alternate method for assessing publication bias in fMRI studies is the relationship between the number of reported foci with the study sample size (e.g., David et al., 2018; David et al., 2013). This test is predicated on the logic that larger studies should be able to detect more differences when the effect is true (David et al., 2018). The values entered into this analysis are reported in the Supplement. There was not a significant relationship between study sample size and the number of reported foci (Pearson r = 0.66, p=.338; Spearman rho=0.40, p=.750), however it is notable that only 4 studies were entered into this analysis.

### 3.4 Discussion

Considering the influence of CRUNCH in the scientific literature (the Google Scholar search identified 283 results; the Reuter-Lorenz & Cappell (2008) manuscript currently shows 813 Google Scholar citations), a surprisingly small number of studies have explicitly tested the model predictions. We attempted to be as inclusive as possible when identifying manuscripts to enter into the analysis. From 52 manuscripts reporting unique results, we identified 9 studies that manipulated task difficulty across three or more levels, compared young to old subjects, and reported the critical behavioural group by load interaction. Of these 9 studies, only 4 reported the critical test of the CRUNCH model predictions for fMRI data. P-curve analysis suggests that while there is evidence to support the CRUNCH model predictions for accuracy data, there is not sufficient evidence to support the predictions for fMRI data.

Importantly, we do not argue that our experimental results (Study 1) or p-curve review (Study 2) invalidate the CRUNCH model or its predictions. Rather, we argue that there is currently insufficient evidence to support (or not support) the CRUNCH model. In other words, we suggest that the CRUNCH in older age literature may show signs of selective reporting, and our experimental results (Study 1) may not be the only results to show inconsistencies with the literature. Overall, existing studies are small (range of n=12-30), and samples are variable (age of subjects in ‘old’ groups vary from 10 to 20 years across studies; estimates of cognitive reserve are inconsistently reported; see Supplement, Study Details Table). Given that the CRUNCH model is currently the most influential model of cognitive compensation in older age, we argue that it is critically important that larger and better controlled studies of the model predictions are conducted.

The scope of the ‘file-drawer problem’ (Rosenthal, 1979), p-hacking (e.g., Head et al., 2015), HARKing (Kerr, 1998), and selective reporting in general, is currently an area of intense interest to researchers. In particular, the fields of experimental psychology and neuroimaging have been shaken by reports of non-reproducibility (Johnson et al., 2017). In a recent survey of 1576 researchers conducted by Nature, more than 70% reported that they had tried and failed to reproduce another scientist’s experiments, and only 23% had attempted to publish a failed reproduction (Baker, 2016). Indeed, when faced with our experimental results (Study 1) – which are in the exact *opposite* direction to the CRUNCH predictions, it was tempting to open the file drawer ourselves. However, we agree with the general conclusion of the literature that cognitive compensation may be a mechanism for maintaining cognitive performance in to older age, and even in the presence of neurodegenerative pathology (e.g., Soloveva et al., 2018). We hope that our results and critical review of the literature will improve our understanding of cognitive compensation, so that ultimately, we can use the evidence to help people maintain healthy cognitive function with high quality of life well into older age.

## 4. Conclusion

Study 1 found no evidence for differential load-dependent changes in fMRI activity in older compared to younger adults, in contrast to the predictions of the CRUNCH model. Study 2 systematically reviewed the CRUNCH literature, and found very few studies that have accurately tested the CRUNCH model predictions. We found evidence for selective reporting in the CRUNCH literature. It is critically important that larger and better controlled studies of the CRUNCH model predictions are conducted.

## Acknowledgements

I thank Gary Egan for valuable discussion of these results, and Nicholas Parsons for assistance with data collection. I thank Richard McIntyre and the research staff of Monash Biomedical Imaging for assistance with data acquisition.

